# A Case Study Leveraging Angler Reported Data For Whirling Disease Monitoring

**DOI:** 10.1101/2025.06.10.658966

**Authors:** Clayton T James, Chloe Christenson, Sean Simmons

**Affiliations:** Angler’s Atlas, Prince George, BC, V2L 4S1; Biological Sciences, University of Alberta, Edmonton, AB, T6G 2E9; Fisheries and Oceans Canada, Edmonton, AB, T6X 0J4

**Author notes:** Ethics and consent statement* - There were no ethical guidelines applicable to this study. Funding statement* – The study was funded independently.

**Keywords:** Alberta, Bow River, citizen science, *Myxobolus cerebralis*, rainbow trout, recruitment, whirling disease

## Abstract

The Bow River (Alberta, Canada) has a well-documented decline in its rainbow trout fishery, with several stressors attributed to this decline including whirling disease (WD). This study evaluated whether anglers could detect recruitment failure in WD-impacted areas of the Bow River drainage using a smartphone app-based citizen science initiative. Anglers reported age 1 and 2 rainbow and cutthroat trout identified by size distribution, estimated from a provincial database, across 4 sub-watersheds with varying densities of *Myxobolus cerebralis* actinospores (TAMs), a known predictor of WD severity. TAM densities ranged from undetectable (0 TAMs/L - Waiparous Creek) to moderate (< 0.01 TAMs/L - Sheep and Highwood Rivers) and high (> 0.01 TAMs/L - Jumpingpound Creek). No fish were captured in Jumpingpound Creek over 20.2 angling hours. The Highwood and Sheep Rivers had catch rates of 0.4 and 0.2 age 1 and 2 trout per hour over 30.6 and 27.7 hours, respectively. Waiparous Creek, where WD was absent, had a significantly higher catch rate of 2.2 trout per hour over 20.3 hours. These findings suggest that self-reported angler data can potentially help identify recruitment losses in WD-impacted areas, demonstrating a novel application of citizen science in fisheries research.

## Introduction

Infectious fish diseases can play significant roles in population-level declines, but are often overlooked in fisheries management due to ecosystem complexity and limitations in broad characterizations of disease in wild populations (Marcogliese 2004; Chapman et al 2021). Understanding disease impacts in the context of wild populations is critical for effective management strategies (Cooke et al 2016; Cooke et al 2017). Consequently, practical field-based approaches are needed to quantify disease outbreaks in wild fish populations (Chapman et al 2021).

Whirling disease (WD) has led to severe declines in rainbow trout (*Oncorhynchus mykiss*) populations in the Rocky Mountain region, including Colorado, Montana, and, more recently, Canada (Nehring and Walker 1996; Vincent 1996; James et al 2021). WD is caused by *Myxobolus cerebralis*, a parasite which requires both a salmonid host and the oligochaete worm *Tubifex tubifex* to complete its lifecycle (Hedrick et al 1998; Gilbert and Granath 2003). Two parasitic spore stages complete the life-cycle of *M. cerebralis* and include an immobile myxospore stage, infectious to the tubificid worms, and a free-floating actinospore stage or triactinomyxon (TAM), infectious to the fish host (Hedrick et al 1998; Gilbert and Granath 2003). Acute WD symptoms develop when trout fry are exposed to high TAM densities shortly after emerging from spawning gravel, up to approximately 9 weeks of age (Ryce et al 2005; Elwell et al 2009). WD in wild populations can be quantified by measuring the density of TAMs in water samples (Nehring and Thompson 2003). TAM densities exceeding 0.01 TAMs/L correlate with trout population declines exceeding 90% (Nehring and Thompson 2003; James et al 2021).

WD-driven mortality is often delayed, with symptoms appearing 20 to 140 days post-infection (Ryce et al 2005; Elwell et al 2009). If WD becomes epizootic in a population, impacts are most obvious by the disappearance of age 1 and 2 trout as age 0 survival becomes impaired (Nehring and Walker 1996; Elwell et al 2009). Nehring and Walker (1996) first discovered WD in the Colorado River in 1993 when sampling efforts revealed a near complete collapse of age 1 and 2 rainbow trout, representing only 0.5 and 0.7% of the total catch, respectively.

In 2016, Canada had its first detection of *M. cerebralis* in Alberta (AEP 2018) and the parasite has since been confirmed through much of the southern portion of the province (AEP 2019). The first evidence of WD impacting rainbow trout populations, including a near absence of juvenile trout was found in 2019 in the Crowsnest River, Alberta (James et al 2021). The nearby Bow River, a highly valued socio-economic trout fishery located north of the Crowsnest River has a well-documented rainbow trout decline (Cahill et al 2018) and the conditions necessary for WD to become epizootic (AEP 2019; Barry et al 2021). While angling mortality has been cited as a primary driver of decline, the impact of WD remains poorly understood (Cahill et al 2018; Christensen et al 2021).

Citizen science provides a cost-effective method for monitoring fisheries over large spatial scales compared to traditional surveys (Jackel et al 2021; Johnston et al 2022). Anglers represent a widely available resource for data collection, and mobile app technology has enhanced their potential contribution (Gundelund et al 2021). Since age 1 and 2 trout fall within a catchable size range, angler reported data may be able to support recruitment failure in the wild linked to WD. Recent data suggests multiple Bow River sub-watersheds have TAM densities capable of causing trout population declines (Christenson 2021). This study aimed to determine whether self-reported angler catch data for age 1 and 2 rainbow and cutthroat trout (*Oncorhynchus clarkii*) in 4 Bow River sub-watersheds with varying TAM densities could confirm recruitment failure linked to WD. Both trout species were included due to hybridization in the Bow River (Rasmussen et al 2010; Sinnatamby et al 2019) and their shared susceptibility to WD (Nehring 2010; Sarker et al 2015).

## Methods

### Site selection

Four sub-watersheds, each comprised of several HUC 10 (Hydrologic Unit Code 10) level watersheds, of the Bow River were selected for this study: Jumpingpound Creek, the Sheep River, the Highwood River, and Waiparous Creek (Figure 1). Each sub-watershed has previously been assessed for WD and is considered important habitat for rainbow trout and/or cutthroat trout (Rhodes 2005; Fitzsimmons 2008). WD was determined by estimating the average density of TAMs collected using a filtration technique developed by Nehring and Thompson (2003). TAM filtrations were conducted in each watershed between May and August 2021 as part of a graduate thesis project (Christenson 2021). TAM densities in each watershed averaged 0 TAMs/L (Waiparous Creek - Control), 1.0 × 10^−4^ TAMs/L (Highwood River), 7.8. x 10^−5^ TAMs/L (Sheep River), and 0.02 TAMs/L (Jumpingpound Creek). Waiparous Creek watershed was chosen to serve as a control as no TAMs were detected prior to this study.

**Figure 1.**
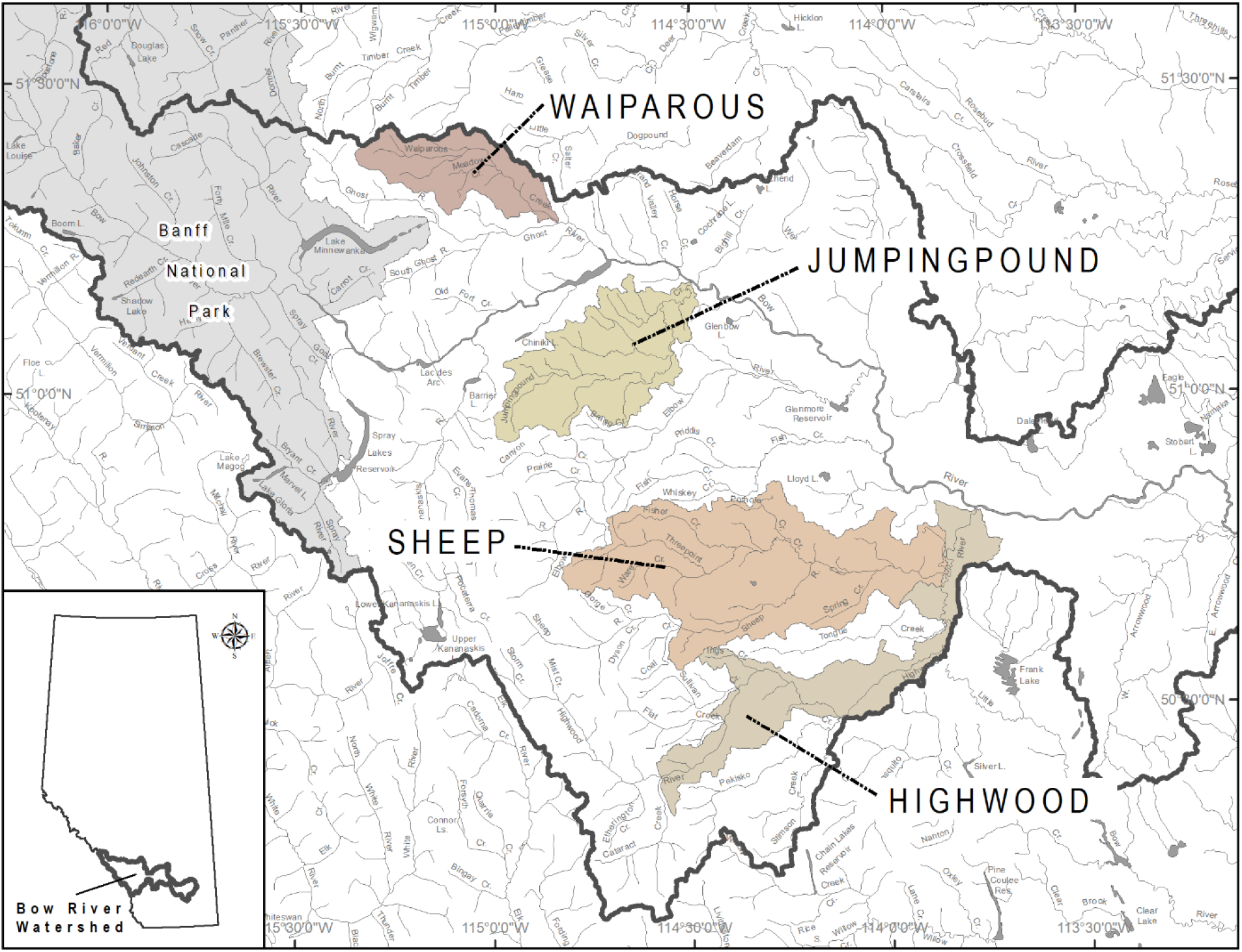
Sub-watershed boundaries (comprised of several HUC 10 watersheds) of Waiparous Creek, the Sheep River, the Highwood River, and Jumpingpound Creek used in this study within the greater Bow River Drainage Basin, AB, Canada.

To demonstrate the presence of sub-adult rainbow trout and cutthroat trout in each sub-watershed prior to the establishment of *M. cerebralis*, catch-per-unit-effort (CPUE) data from previous backpack electrofishing surveys were retrieved from the FWMIS database (AEP 2024). Only data collected before 2016 were included, as 2015 marked the last known year before *M. cerebralis* was confirmed in the Bow River basin.

### App-based event model

Anglers were recruited to collect catch data in each watershed using the MyCatch mobile phone app. Anglers were required to make a profile in the MyCatch app and then register for the event titled ‘*Bow River Rainbow Trout Recruitment Survey*’. For the purposes of this study, anglers were encouraged to capture and report smaller trout within each of the sub-watersheds identified, however, all trout sizes were reported.

Georeferenced boundaries were created for each sub-watershed in this study and anglers were required to have their GPS/location services enabled on their phone for catches to qualify. Self-reported data collection began on June 16, 2023 coinciding with provincial fisheries regulations, and closed September 30, 2023. For each angling trip, anglers reported the date, waterbody, hours fished, and any fish captured identified to species. Fish captured also required a total length measurement with a photograph taken through the MyCatch app. Photographs and measurements were screened and verified by MyCatch staff after catch and trip data had been synchronized with online servers.

### Statistical analyses

A length-frequency distribution for rainbow trout and cutthroat trout in the Bow River was created using data from the provincial fisheries database (FWMIS) (AEP 2024) to determine the typical fork length range of age 1 and age 2-year classes (herein referred to as sub-adults). Based on this, sub-adult trout were defined as those between 80 and 240 mm fork length.

Differences in angling catch rates of sub-adult rainbow trout and cutthroat trout between sub-watersheds were assessed using a Kruskal-Wallis rank sum test, as the data did not meet the assumptions of normality or equal variances. Pairwise comparisons were made using the Wilcoxon rank sum test. A generalized linear model was applied to assess the relationship between sub-adult trout catch rate and TAM density. TAM density data were log-transformed to meet normality requirements, as well a correction factor of 0.0000000001 was added to all TAM averages before log transformation to avoid having a 0 in the data. All statistical analyses were performed using R version 4.2.2 (2022).

## Results

### Historical Electrofishing CPUE

Historical backpack electrofishing catch-per-unit-effort (CPUE) data from 1997 to 2015, summarized in Table 1, indicated that each sub-watershed previously supported a stable population of sub-adult trout prior to the establishment of *M. cerebralis* in the Bow River basin. Each sub-watershed sampled had a minimum of 3 years of data, with average CPUEs for sub-adult rainbow trout and cutthroat trout ranging from 0.27 to 0.91 fish per 100 seconds of electrofishing effort.

**Table 1.** Backpack electrofishing CPUEs (fish / 100 sec) of sub-adult rainbow trout and cutthroat trout from 1997 to 2015 in four sub-watersheds of the Bow River.

| Sub-watershed | Years sampled | Sampling events (n) | Avg CPUE | CPUE range |
| --- | --- | --- | --- | --- |
| Waiparous Creek (Control) | 2000, 2003, 2006, 2010, 2011, 2015 | 57 | 0.27 | 0.02 – 1.01 |
| Highwood River | 1998, 2006, 2015 | 5 | 0.91 | 0.35 – 2.02 |
| Sheep River | 1997, 1998, 2005, 2006, 2007, 2010, 2011, 2014 | 38 | 0.64 | 0.05 – 2.52 |
| Jumpingpound Creek | 1997, 1998, 2003, 2005, 2006, 2007, 2008, 2011, 2015 | 53 | 0.62 | 0.04 – 6.36 |

### App-based angling catch rates

A total of 26 anglers registered for the ‘*Bow River Rainbow Trout Recruitment Survey*’ event in 2023; however, only 8 anglers contributed effort and catch data via the MyCatch app. Over the course of the study, 72 fish were captured, of which 48 were sub-adult rainbow trout and cutthroat trout. Angling effort ranged from 20.2 to 30.6 hours per watershed, with average trip lengths between 2.2 and 3.8 hours. Catch rates for sub-adult trout varied between sub-watersheds. In Waiparous Creek (control), the catch rate averaged 2.2 fish per hour, which was significantly higher than in any other sub-watershed (*P* < 0.001) (Figure 2). The catch rates for sub-adult trout in the Highwood River and Sheep River were similar, with average rates of 0.4 and 0.2 fish per hour, respectively, and were not significantly different (*P* = 0.3) (Figure 2). No fish of any size were reported in Jumpingpound Creek despite 20.2 hours of angling effort.

**Figure 2.**
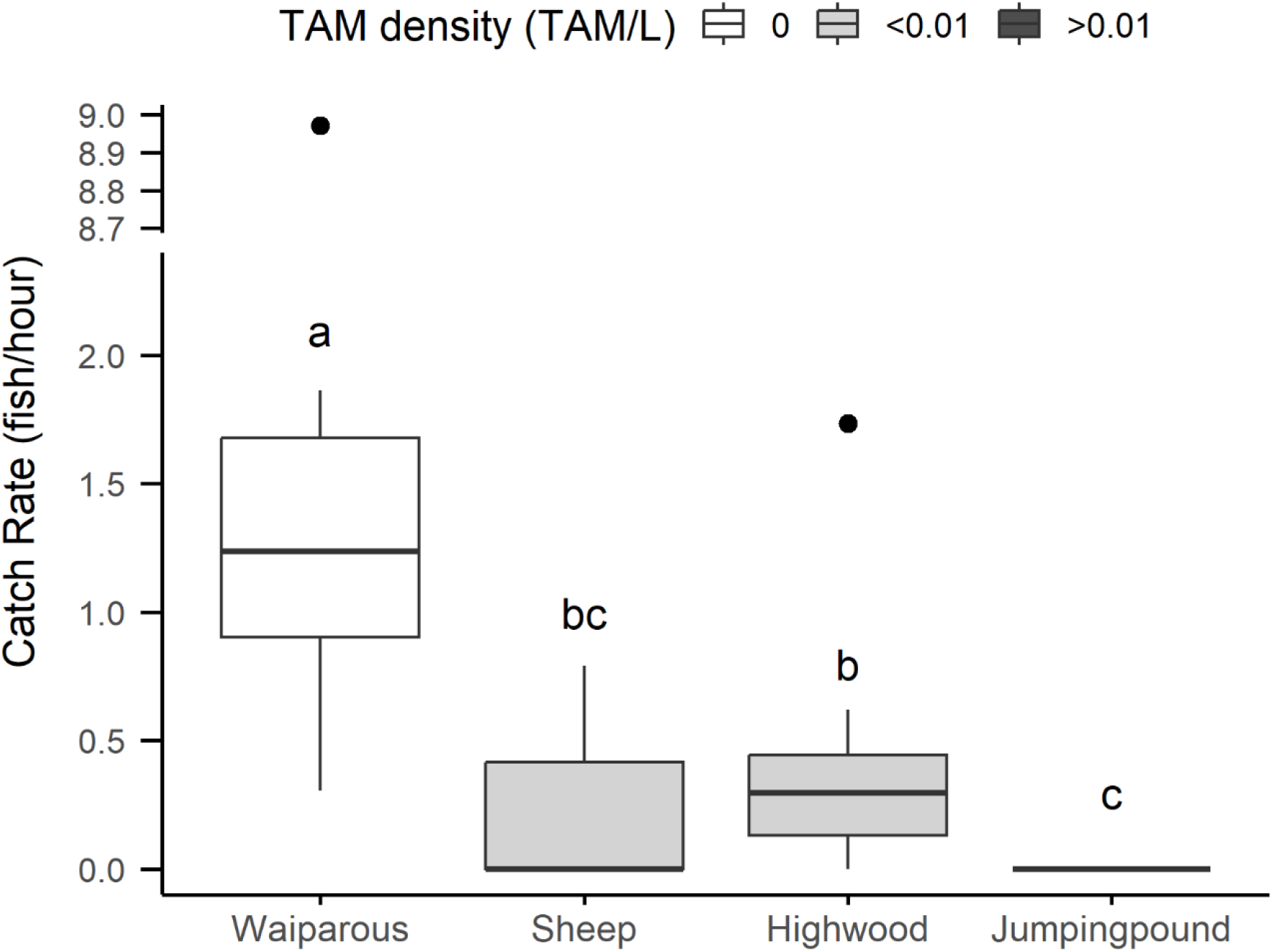
Box plots of angling catch rates for Waiparous Creek, Sheep River, Highwood River, and Jumpingpound Creek between June 16 and September 30, 2023. Sites are distinguished by previous TAM density measurements (0 TAMs/L, <0.01 TAMs/L, and >0.01 TAMs/L). Lower and upper fences are 25th and 75th percentiles, and the median is denoted as the line within each box. Bars represent 10th and 90th percentiles and letters above each bow denote significance. Outliers are denoted as a dot falling outside the 10th and 90th percentiles.

### Relationship between catch rate and TAM density

Despite the small sample size of sub-watersheds (n = 4), a significant negative relationship between angling catch rate and TAM density (log transformed) was observed (*R* = 0.95, *P* = 0.03) (Figure 3).

**Figure 3.**
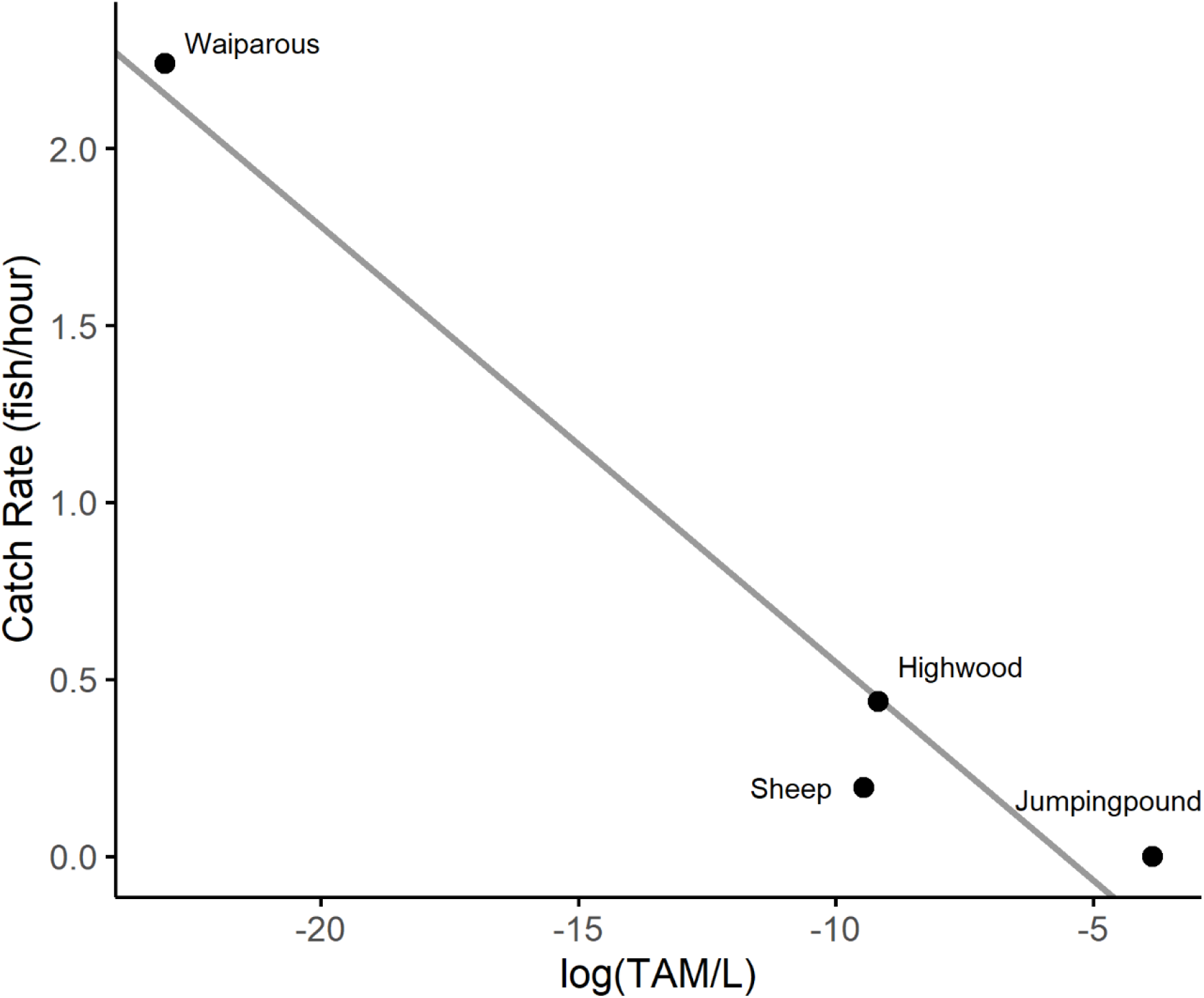
Scatter plot showing the relationship between log transformed TAM density (TAMs/L) and catch rate (sub-adult fish/hr) for 4 sub-watersheds of the Bow River. The linear regression line (solid grey line) described by the equation Catch rate=-0.68-0.12*log(TAM) represents the best-fit model explaining the variation in catch rate based on TAMs/L. The model accounts for 95% of the variation in catch rate.

## Discussion

This study presents a novel approach to using self-reported angler catch data to identify possible recruitment failure in fish populations affected by whirling disease. Our findings indicate that trout recruitment is significantly impaired in sub-watersheds with moderate to high densities of *Myxobolus cerebralis* actinospores (TAMs). The absence of fish of any size captured in Jumpingpound Creek, which exceeded the known TAM density threshold for severe population declines, strongly suggests an epizootic-level WD outbreak. This aligns with historical data confirming Jumpingpound Creek as a formerly productive trout fishery (Jumpingpound Creek Watershed Partnership, 2009) and highlights the need for further investigation.

Catch rates of sub-adult trout in the Highwood and Sheep River sub-watersheds were also low, supporting the observed negative relationship between WD prevalence and recruitment. Despite these sub-watersheds having TAM densities below the 0.01 TAMs/L threshold, our results suggest even moderate detectable densities of TAMs in the water may still impair recruitment. Meanwhile, Waiparous Creek, where TAMs were undetected, exhibited significantly higher catch rates, reinforcing the correlation between TAM density and recruitment loss. Waiparous Creek has been confirmed for the establishment of *M. cerebralis* during previous molecular testing in 2018 (AEP 2019). This indicates that the ecological/environmental conditions of Waiparous Creek during these studies currently may preclude *M. cerebralis* from reaching an epizootic status as no TAM detections were found. Reasons may include low densities of *T. tubifex* worms or inadequate temperatures to support TAM proliferation. Further investigation and monitoring in Waiparous Creek are recommended in order to inform future risk potential of WD outbreak.

Despite clear evidence of *Myxobolus cerebralis* parasitic amplification in both hosts of its two-host life cycle, misconceptions persist among some anglers and even biologists, leading to an underestimation of its impact on trout populations. Once the parasite becomes enzootic, extreme levels of amplification occur in both the fish-to-worm and worm-to-fish stages. Nehring et al (2014) demonstrated that a single cohort of *T. tubifex* worms exposed to 12,500 myxospores produced over 22 million TAMs over 180 days, with each TAM containing at least 32 to 64 infective sporoplasms, exponentially increasing infection potential. Similarly, El-Matbouli and Hoffmann (1998) showed that a single ingested myxospore undergoes up to 15 rounds of division in the worm’s gut, drastically increasing TAM output. This extreme amplification results in high mortality when trout fry encounter even low TAM doses. Nehring et al (2003) found that just 3 to 5 TAMs per fry over two weeks led to 10.44% mortality, while exposure to 286 to 404 TAMs resulted in 66.15% mortality over 180 days. Given these findings, dismissing whirling disease as a primary driver of trout recruitment collapse, particularly in areas with detectable levels of TAMs in the water, reflects a lack of awareness of the parasite’s biological dynamics in natural systems.

Citizen science in fisheries research is often limited to broad monitoring, such as creel surveys, but this study demonstrates it has potential for detecting specific ecological patterns, including recruitment failure due to disease. However, low angler participation and a small sample size were limitations. While 26 anglers registered, only 8 contributed data across 34 trips. Low engagement is a common challenge in citizen science (Venturelli et al 2017), often attributed to limited awareness of research objectives and competing recreational interests (Crandall et al 2018). Our initial outreach to local fishing clubs had limited success, and social media campaigns launched mid-study may have been too delayed to attract broader participation. Additionally, asking anglers to target small fish may have reduced engagement, as larger fish are typically more desirable catches. Despite these challenges, effort was relatively consistent across sub-watersheds, suggesting comparable fishing conditions.

While preliminary, these findings indicate that anglers can serve as effective, cost-efficient monitors of recruitment failure linked to WD. Although angler-reported observations are subject to biases and are limited in this study, they align with and help contextualize ecological survey data, particularly the observed density of TAMs, which remains the clearest signal of WD impact. In this way, angler reports augment, rather than replace, traditional monitoring efforts. Further research is needed to quantify the full extent of WD impacts in the Bow River system. Jumpingpound Creek, in particular, warrants immediate attention due to the apparent collapse of its trout population. Although the Highwood and Sheep Rivers have lower TAM densities, ongoing monitoring is necessary to assess whether WD-related impacts will intensify over time. Without addressing these data gaps, fisheries managers risk implementing ineffective or even counterproductive regulations. For instance, if WD is a primary driver of trout declines, restricting angling effort will not mitigate its impact and may instead redirect anglers to other vulnerable fisheries. The Bow River alone had nearly 200,000 angling hours between June and September 2018 (Christensen et al 2021), and displacing this effort could strain smaller fisheries or facilitate the spread of *M. cerebralis* to currently uninfected waters.

The Bow River remains a highly valued fishery, with many angling groups actively engaged in its conservation. Overlooking the contributions anglers can make to fisheries management undermines conservation efforts. When initial electrofishing studies on the Upper Colorado in October 1993 revealed the loss of age 1 and 2 cohorts of rainbow trout (Nehring and Walker 1996), notably, experienced fly fishermen, including Charlie Meyers, a longtime outdoor writer for the *Denver Post*, reported a noticeable absence of juvenile rainbow trout (6 to 8 inches) in their catches as early as 1991 and 1992 (R Barry Nehring, Colorado Parks and Wildlife, Fort Collins, CO, pers. comm.). According to longtime Bow River angler Greg Allard (pers. comm.), there has been a steady decline in juvenile rainbow trout abundance over the last 20 years. Once-common sightings of young-of-year trout using weedy nursery banks for refuge and large schools of 1-year-olds returning near the Fish Creek confluence in late summer have become rare or greatly diminished. Allard notes that many veteran Bow River anglers have observed similar declines over the past 2 decades. This emphasizes that dedicated anglers can provide valuable early insights into concerning ecological changes, often recognizing trends before biologists and scientists. Integrating anglers into future monitoring initiatives could enhance data collection efforts and promote broader support for fisheries research and management. This study highlights the potential of citizen science to fill critical knowledge gaps in fish disease ecology and underscores the need for continued research on WD impacts in Alberta’s trout fisheries.

## Acknowledgments

We thank Barry Nehring, Clayton Raines, Brandi Mogge and Tyana Rudolfsen for their comments on previous versions of this manuscript. In addition, we thank David Blair and the Bow River Trout Foundation for encouraging angling groups to contribute data to this study. These groups include the Calgary Fish and Game, Calgary Women Fly Fishers Club, Sarcee Fish and Game, Angling and Outfitters, and Hook and Hackle Club. We give specific thanks to the following anglers for assisting in data collection: Jill Collyer, Kyle Bruce, Rod Day, Mike Duszynski, Brett Husband, and Laurence Mattson. We also thank the MyCatch Team, specifically Miyah Clarke, Jim Clarke, and Nadine Gordon for successful implementation of this event in the MyCatch app.

